# A selective cortico-limbic network organizes behavior during reward seeking under threat

**DOI:** 10.64898/2026.03.12.711336

**Authors:** Rodrigo O. Sierra, Leticia Ramirez-Lugo, Elizabeth Illescas-Huerta, Axel R. Bolaños, Francisco Sotres-Bayon

## Abstract

To obtain rewards, animals must select actions while facing threats, often under competing appetitive and defensive drives and with uncertainty about harm. While neural circuits controlling isolated threats or rewards are well characterized, it remains unclear which specific cortical and subcortical nodes are causally necessary to organize behavior during motivational conflict. Here, we used the step-down avoidance-mediated conflict (SDAmC) task in adult male rats, which quantifies avoidance, risk assessment, and reward approach within the same session. We performed pharmacological inactivation across eight candidate structures implicated in valence and action selection: prelimbic (PL) and infralimbic (IL) cortices, lateral orbitofrontal cortex (lOFC), anterior (aIC) and posterior (pIC) insular cortices, lateral habenula (LHb), basolateral amygdala (BLA), and nucleus accumbens (NAc). Inactivation revealed a precise anatomical dissociation within this network. Silencing PL or pIC facilitated approach behavior during conflict, but with dissociable effects on risk assessment. In contrast, BLA inactivation induced a broader behavioral disinhibition evident even in non-conflict conditions, whereas NAc inactivation disrupted the temporal organization of approach, yielding a fragmented behavioral phenotype. Notably, inactivation of IL, lOFC, aIC, and LHb did not alter conflict resolution in this paradigm. Together, these findings identify a selective cortico-limbic network in which PL and pIC regulate conflict-specific approach, BLA maintains defensive thresholding, and NAc coordinates goal-directed action, constraining the set of brain regions that are necessary to organize behavior when reward seeking competes with threat.

**Significance Statement:** In threatening contexts, animals must resolve motivational conflict by integrating defensive and appetitive drives. Although threat and reward circuits have been well studied, the neural nodes that organize avoidance, risk assessment, and approach during conflict remain poorly defined. Here, we used a validated conflict paradigm and pharmacological inactivation to test multiple cortical and subcortical candidates. Motivational conflict depends on a restricted subset of structures: the prelimbic cortex and posterior insula selectively gate approach, the basolateral amygdala supports defensive constraint, and the nucleus accumbens coordinates goal-directed action. Other regions were dispensable. This study provides causal evidence for a constrained functional architecture underlying motivated decisions under threat.

## Introduction

Animals routinely face situations in which approaching a reward requires entering a potentially harmful context. Such approach-avoidance conflicts constitute a distinct form of decision making in which appetitive and defensive motivational systems, supported by partially distinct neural circuits for threat processing (LeDoux, 2000; Maren, 2001; Tovote, Fadok, & Luthi, 2015) and reward seeking (Berridge & Robinson, 1998; Everitt & Robbins, 2005; Floresco, 2015), must be integrated to select a single course of action. Classical behavioral theory formalized this dilemma as competing approach and avoidance gradients (Miller, 1944). Subsequent conflict paradigms operationalized it by placing reward seeking under the threat of punishment, such as punished responding and punished drinking procedures (Paterson & Hanania, 2010; Vogel, Beer, & Clody, 1971). Across species and behavioral tasks, conflict is expressed not simply as suppression of approach, but as a structured sequence of behaviors that includes hesitation, risk assessment, and action commitment. Sensitivity to anxiolytic manipulations highlights the translational relevance of these behaviors (McNaughton & Corr, 2014). Importantly, approach-avoidance conflict is not reducible to threat or reward alone but emerges when competing motivational values must be resolved under uncertainty (Calhoon & Tye, 2015; Tye, 2018).

Despite extensive research on the neural circuits underlying threat and reward, it remains unclear how specific cortical and subcortical nodes contribute to the internal organization of behavior during conflict. Medial prefrontal regions, including the prelimbic and infralimbic cortices, regulate defensive expression and its suppression across threat-learning and avoidance paradigms (Bravo-Rivera, Roman-Ortiz, Brignoni-Perez, Sotres-Bayon, & Quirk, 2014; Sotres-Bayon & Quirk, 2010; Vidal-Gonzalez, Vidal-Gonzalez, Rauch, & Quirk, 2006), whereas the orbitofrontal cortex supports value-based action selection when outcomes involve competing costs and benefits (Orsini, Moorman, Young, Setlow, & Floresco, 2015). The insular cortex links the interoceptive state to aversive valuation and motivational drive (Gehrlach et al., 2019), and subcortical structures, such as the basolateral amygdala and nucleus accumbens, play central roles in assigning motivational significance and guiding goal-directed behavior (Stuber et al., 2011). In addition, the lateral habenula has been implicated in negative value signalling and aversive bias during decision making under uncertainty (Shabel, Proulx, Piriz, & Malinow, 2014). However, much of this literature derives from tasks that isolate threat or reward, leaving unresolved which of these regions are causally required to organize the sequence and coupling of avoidance, risk assessment, and approach that define motivated conflict. Establishing node-specific necessity, including both positive and negative contributions, is therefore critical for constraining circuit-level models of conflict behavior (Choi, Jean-Richard-Dit-Bressel, Clifford, & McNally, 2019; Friedman et al., 2015).

To address this issue, we used the step-down avoidance–mediated conflict (SDAmC) task, a paradigm that dissociates avoidance, risk assessment, and approach within a single behavioral episode and has been previously validated pharmacologically (Illescas-Huerta, Ramirez-Lugo, Sierra, Quillfeldt, & Sotres-Bayon, 2021; Pereira, Ardenghi, Mello e Souza, Medina, & Izquierdo, 2001; Perry, Dias, Carrasco, & Izquierdo, 1983). In SDAmC, rats must decide whether to descend from a safe platform onto a shock-associated grid to obtain a palatable reward, allowing a direct comparison between conflict and non-conflict conditions by manipulating motivational state (thirsty or not). This design isolates behavioral components that are genuinely conflict-dependent rather than reflecting reward seeking or threat memory alone. To establish causal necessity at the regional level, we used pharmacological inactivation, examining its contribution across multiple cortical and subcortical regions implicated in threat, valuation, interoception, and motivation, including the prelimbic and infralimbic cortices, lateral orbitofrontal cortex, anterior and posterior insular cortex, lateral habenula, basolateral amygdala, and nucleus accumbens (Bravo-Rivera et al., 2014; Floresco, 2015; Gremel & Costa, 2013; Moorman & Aston-Jones, 2015; Parkes, Bradfield, & Balleine, 2015; Ramirez-Lugo, Penas-Rincon, Angeles-Duran, & Sotres-Bayon, 2016; Stopper & Floresco, 2014; Velazquez-Hernandez & Sotres-Bayon, 2021; Wassum & Izquierdo, 2015). By quantifying both the magnitude and the internal organization of behavioral components, we identify a selective subset of regions that exert dissociable control over risk assessment and action selection during conflict. These findings define a constrained functional architecture in which selective subset of cortico-limbic hubs organize behavior when reward seeking occurs under threat.

## Materials and Methods

### Subjects

All experimental procedures were approved by the Institutional Animal Care and Use Committee of the Universidad Nacional Autónoma de México (UNAM) and were conducted in accordance with the National Ministry of Health guidelines for the care and use of laboratory animals. A total of 208 adult male Wistar rats (Instituto de Fisiología Celular, UNAM breeding colony), 2–3 months old and weighing 250–300 g at the start of the experiments, were used. Rats were housed individually in polyethylene cages to monitor water consumption, handled daily, and maintained on a 12-h light/dark cycle. All experiments were performed during the light phase. To maintain stable motivation to seek water, rats assigned to the conflict condition were water-restricted to 12 mL/day (6 mL in the morning and 6 mL in the afternoon). Rats assigned to the non-conflict condition had *ad libitum* access to water, except during specific training phases, as described below, to ensure task acquisition.

### Surgery

Rats were anesthetized with isoflurane (5% induction, 2–3% maintenance) and placed in a stereotaxic frame (Kopf Instruments). Stainless-steel guide cannulae (23-gauge) were bilaterally implanted targeting the prelimbic cortex (PL; AP +3.0, ML ±0.6, DV −3.0), infralimbic cortex (IL; AP +3.0, ML ±3.1, DV −3.8; 30° angle), anterior insular cortex (aIC; AP +2.7, ML ±3.9, DV −5.2), posterior insular cortex (pIC; AP −1.8, ML ±6.2, DV −6.2), lateral orbitofrontal cortex (lOFC; AP +3.7, ML ±2.6, DV −3.6), basolateral amygdala (BLA; AP −2.8, ML ±5.0, DV −7.4), nucleus accumbens (NAc; AP +2.5, ML ±1.5, DV −5.8), and lateral habenula (LHb; AP −3.6, ML ±0.7, DV −4.0) (Paxinos et al., 1998). Coordinates are expressed in millimeters relative to bregma. Cannulae were secured with stainless-steel screws and dental acrylic, and obturators were inserted to maintain patency until microinfusions. Rats were allowed at least 7 days of recovery prior to behavioral procedures.

### Step-down avoidance-mediated conflict (SDAmC) task and behavioral protocol

Behavioral testing was conducted in a modified step-down avoidance chamber (50 × 25 × 25 cm) housed within a sound-attenuating cubicle (**Fig. 1A**). The apparatus consisted of a “Safe Zone” (elevated acrylic platform; 20 × 23.5 × 6 cm) and a “Threat Zone” (stainless-steel grid floor; 30 × 25 cm) connected to a shock generator (Coulbourn Instruments). A removable acrylic sliding door separated the platform from the grid floor. On the wall opposite to the platform, a bottle containing 0.1% saccharin served as the reward. Between sessions, the apparatus (grid, floor, and walls) was cleaned with soapy water and 70% alcohol, then dried with paper towels. For behavioral quantification, the apparatus was divided into a start zone (platform), a choice zone (proximal grid adjacent to the platform), and a reward zone (distal grid containing the saccharin bottle).

**Figure 1.**
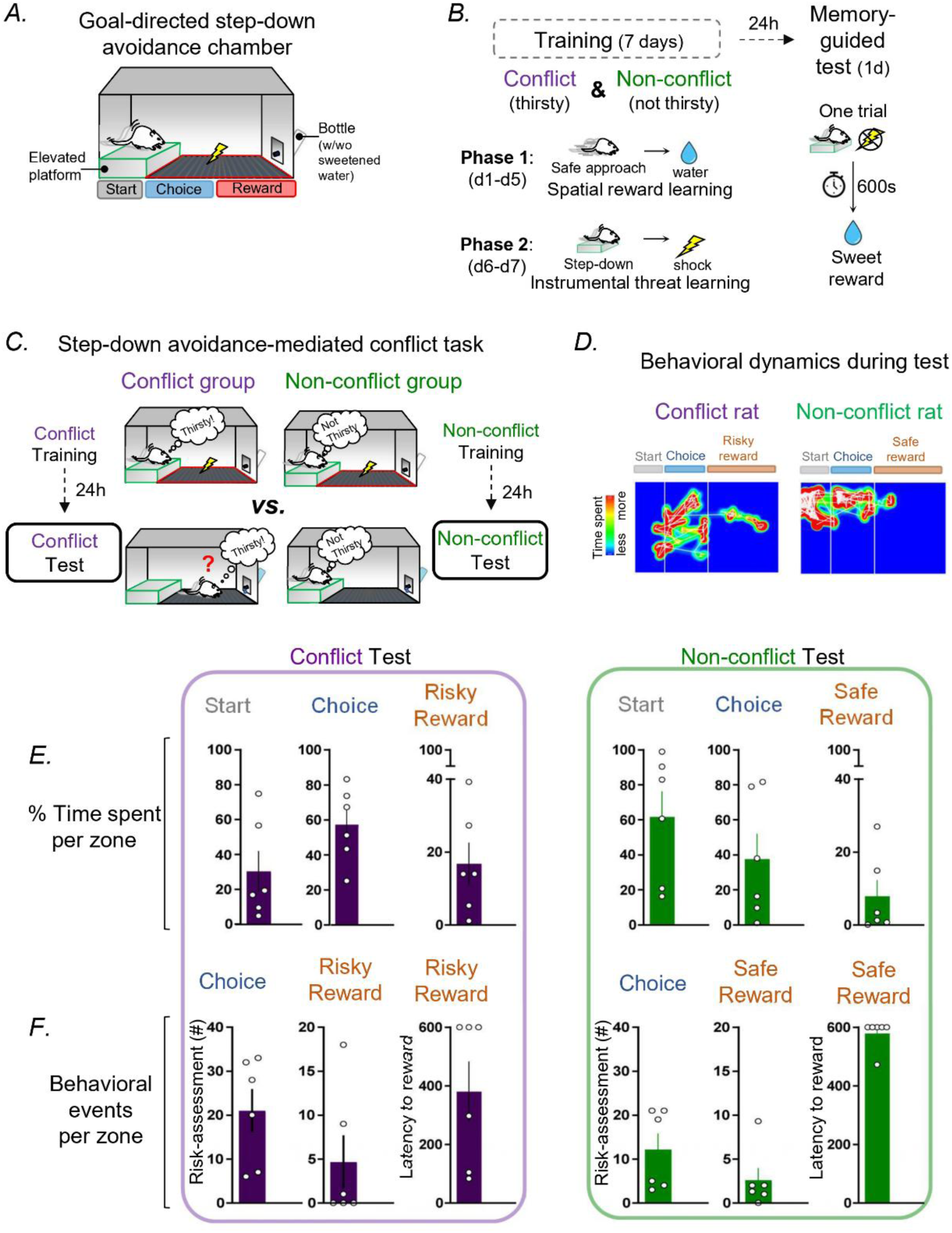
Step-down avoidance–mediated conflict (SDAmC) task and behavioral measures. (A) Schematic of the SDAmC apparatus showing the start zone (elevated platform), choice zone (proximal grid), and reward zone (distal grid containing 0.1% saccharin). (B) Experimental design and timeline. Rats underwent reward conditioning in Phase 1 (Days 1–5), followed by threat conditioning in Phase 2 (Days 6–7; 0.5 mA shock delivered upon stepping down and terminated upon return to the platform). On the test day (Day 8), animals were tested without shock for up to 600 s under either conflict (water-restricted; thirsty) or non-conflict (sated) conditions. (C) Test conditions highlighting the motivational manipulation that distinguishes conflict (risky reward) from non-conflict (safe reward) performance. (D) Representative movement heatmaps illustrating spatial occupancy during test sessions. (E–F) Quantification of time spent in each zone (E) and behavioral measures, including step-down latency and risk assessment events (F). Conflict group, n = 6; non-conflict group, n = 6. Data are expressed as mean ± SEM. Individual data points are represented by open circles.

The protocol was conducted across 10 consecutive days and consisted of habituation, reward conditioning, threat conditioning, and a conflict test (**Fig. 1B**). During habituation (two sessions), water-restricted rats were placed on the platform for 5 min with the door closed, after which the door was opened and animals explored the apparatus for 10 min; no shock or reward was delivered. During Phase 1 (reward conditioning, Days 1–5), water-restricted rats were placed on the platform and, after 5 min, were allowed access to saccharin; step-down latencies were recorded daily. During Phase 2 (threat conditioning, Days 6–7), rats maintained on the water-restriction schedule were placed on the platform; upon stepping down onto the grid with all four paws, a mild footshock (0.5 mA) was delivered continuously until the rat returned to the platform. On the test day (Day 8), no shocks were delivered, and sessions lasted up to 600 s. Conflict animals were tested under water restriction (thirsty), creating conflict between reward approach and avoidance of the shock-associated grid. Non-conflict animals were provided *ad libitum* water for 24 h prior to testing (sated), minimizing appetitive motivation.

### Drug infusion

Bilateral microinfusions were performed 15 min prior to the test session (Day 8) or the saccharin intake test. To inactivate target regions, a cocktail of GABA receptor agonists (250 ng/µL + baclofen 250 ng/µL; Sigma-Aldrich) dissolved in sterile saline was infused, while control animals received saline vehicle. Infusions were delivered through 30-gauge injectors extending 1 mm beyond the guide cannulae and connected to 10 µL Hamilton syringes via polyethylene tubing. A total volume of 0.3–0.5 µL per hemisphere (depending on target region) was infused at 0.4 µL/min using a microinfusion pump. Injectors were left in place for an additional 1 min to allow diffusion.

### Saccharin intake test

To control for nonspecific effects on fluid consumption, an independent sample of water-restricted rats was tested in the home cage. Animals were presented with 0.1% saccharin for 30 min, and consumption was quantified by bottle weight before and after the session.

### Histology

After behavioral testing, rats were deeply anesthetized with an overdose of pentobarbital and perfused transcardially with 0.9% saline followed by 4% paraformaldehyde. Brains were removed, post-fixed, and cryoprotected in 30% sucrose. Coronal sections (40 µm) were cut and Nissl-stained (cresyl violet) to verify cannula placements. Only data from animals with injector tips located within the intended target boundaries were included in statistical analyses.

### Data collection and behavioral analysis

Behavior was recorded with an overhead digital video camera and analyzed using ANY-maze (Stoelting). Automated tracking used three body points (head, center, tail). Zone occupancy was determined using the center point for start and reward zones and the head point for the choice zone (entry threshold: 10 mm) to register risk assessment-related probes. Step-down latency was defined as the time from door opening to placement of all four paws on the grid (choice zone entry). Time spent in start, choice, and reward zones was quantified for each session and expressed as both duration and percentage of total session time. Risk assessment events were defined as body elongation from the platform toward the grid followed by retraction and were quantified using head-point entries ≥10 mm into the choice or reward zone. Movement heatmaps were generated from head coordinates to visualize exploration patterns, with the color scale maximum set to 10 s for peak intensity.

### Statistical Analysis

All analyses were performed in GraphPad Prism 10.0. Data were assessed for normality using the Kolmogorov–Smirnov test. Because behavioral variables did not consistently meet normality assumptions, comparisons between independent groups were conducted using the nonparametric Mann–Whitney U test. To characterize behavioral organization, Spearman rank correlations (rs) were computed for brain regions showing significant effects in the conflict condition. Correlation matrices included six variables: time spent in the start, choice, and reward zones; risk-assessment event frequency in the choice and reward zones; and step-down latency. To compare aggregate behavioral profiles across structures, each behavioral correlation matrix was flattened into a vector, and pairwise correlations were computed between structures. Data are presented as mean ± SEM, and statistical significance was set at p < 0.05.

## Results

Rats were trained and tested on the SDAmC task, which dissociates avoidance, risk assessment, and reward approach using discrete spatial zones (start, choice, reward) within a single session (**Fig. 1A–C**). Motivational state (thirsty vs. sated) produced a robust shift in behavioral allocation and decision dynamics across these readouts, providing a structured framework to quantify how regional inactivation alters conflict performance (**Fig.–F**).

### Prelimbic cortex inactivation biases conflict resolution toward approach

We targeted the prelimbic cortex (PL; cannula placements shown in **Fig. 2A**) given its established involvement in defensive control, persistent threat monitoring, and action selection under uncertainty (Bravo-Rivera et al., 2014; Burgos-Robles, Vidal-Gonzalez, & Quirk, 2009; Capuzzo & Floresco, 2020; Sotres-Bayon, Sierra-Mercado, Pardilla-Delgado, & Quirk, 2012). Representative movement heatmaps illustrate that PL inactivation increased exploration of grid zones during conflict relative to vehicle, with no obvious change under non-conflict conditions (**Fig. 2B**). During the conflict test, vehicle-treated rats displayed a strongly avoidance-biased strategy, remaining predominantly in the start zone with limited transitions onto the grid (**Fig. 2B, top**). In contrast, PL inactivation produced robust and selective changes: time in the start zone decreased (Mann–Whitney, U = 104, p = 0.016; **Fig. 2C**), time in the choice zone increased (Mann–Whitney, U = 26, p = 0.016; **Fig. 2C**), and step-down latency was markedly reduced (Mann–Whitney, U = 17, p < 0.001; **Fig. 2D**), consistent with faster commitment to enter the grid. Notably, risk assessment frequency in both the choice and reward zones was unchanged (**Fig. 2D**), suggesting that PL inactivation facilitated approach without a parallel increase in overt risk sampling.

**Figure 2.**
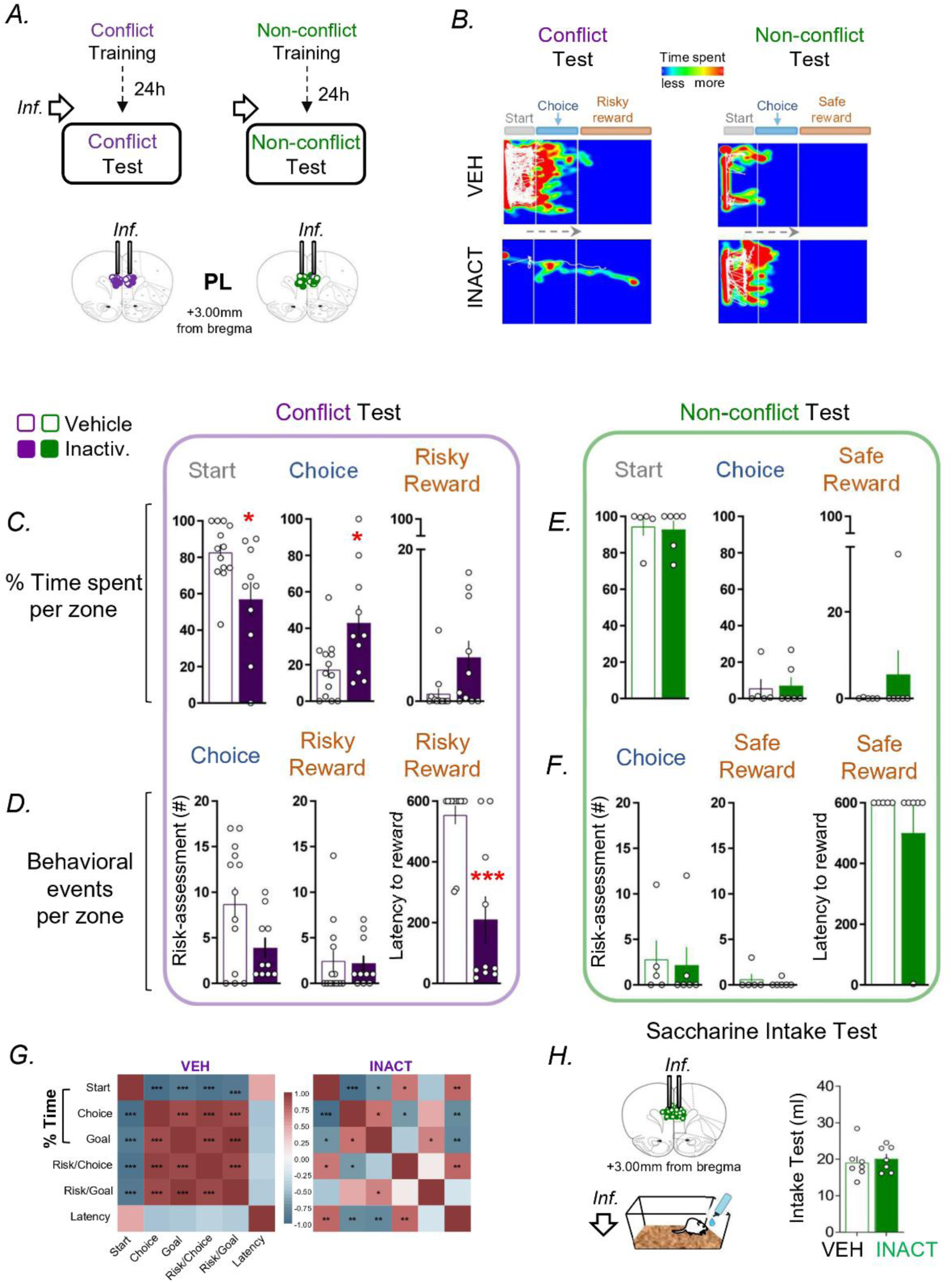
Prelimbic cortex (PL) inactivation selectively facilitates approach during conflict. (A) Cannula placements for PL and experimental timeline indicating microinfusions prior to Conflict or Non-conflict testing. (B) Representative heatmaps during Conflict and Non-conflict sessions under vehicle (VEH) or inactivation (INACT). (C–D) Conflict test quantification showing that PL inactivation reduced time in the start and increased time in the choice zone, with a marked reduction in step-down latency, while risk assessment frequency was unchanged. (E–F) Non-conflict test measures showing no detectable effects of PL inactivation under satiety. (G) Correlation matrices illustrating behavioral organization during Conflict under VEH versus INACT conditions. (H) Home-cage saccharin intake showing no effect of PL inactivation on consummatory behavior (vehicle, n = 7; inactivation, n = 7). Conflict group (vehicle, n = 13; inactivation, n = 10); non-conflict group (vehicle, n = 5; inactivation, n = 5). *p < 0.05; **p < 0.01; ***p < 0.001. Data are shown as mean ± SEM. Individual data points are represented by open circles.

To test whether these effects reflected impaired threat memory retrieval or nonspecific locomotor changes, we examined a separate cohort tested under non-conflict conditions (sated). In the absence of conflict, PL inactivation did not alter any behavioral measure (all p > 0.05; **Fig. 2E–F**), indicating that PL recruitment is selectively engaged when motivational competition is present. Next, we asked whether PL inactivation altered not only behavioral outputs but also their internal coordination. Under vehicle conditions during conflict, behavioral variables showed a coherent structure: time in start was strongly and negatively correlated with time in choice (rs = −1.00, p < 0.001) and reward (rs = −0.89, p < 0.001). Risk assessment events in choice and reward were strongly coupled (rs = 0.92, p < 0.001), and risk assessment in choice was negatively correlated with time in start (rs = −0.96, p < 0.001). This organization was disrupted by PL inactivation: the coupling between risk assessment events in choice and reward was abolished (rs = 0.08, p = 0.828), the relationship between time in start and risk assessment in choice reversed direction (rs = 0.70, p = 0.025), and time in start became positively associated with step-down latency (rs = 0.77, p = 0.010), an association not observed under vehicle conditions (**Fig. 2G**). Finally, saccharin intake in the home cage was unaffected, arguing against nonspecific changes in appetitive motivation (**Fig. 2H**). Overall, PL activity maintains an avoidance-biased strategy during motivational conflict and preserves the coordination linking avoidance, risk sampling, and action initiation.

### Inactivation of IL and lOFC does not alter conflict behavior

To determine whether PL effects reflected a broader frontal cortical contribution, we tested the infralimbic cortex (IL; **Fig. 3A1**), implicated in suppressing defensive responses (Sierra-Mercado, Padilla-Coreano, & Quirk, 2011), and the lateral orbitofrontal cortex (lOFC; **Fig. 3A2**), implicated in value-based updating and action selection (Gremel & Costa, 2013; Ramirez-Lugo et al., 2016). Representative heatmaps show similar occupancy patterns in vehicle and inactivation conditions for both IL and lOFC during conflict testing (**Fig. 3B1–B2**). Consistent with this, inactivation of either region did not alter conflict performance: time allocation across the start, choice, and reward zones, risk assessment frequency, and step-down latency were comparable between vehicle and inactivation conditions (all p > 0.05; **Fig. 3C1–C2, D1–D2**). Because IL and lOFC inactivation produced no detectable effects during conflict, these regions were not examined further in non-conflict, saccharin intake, or correlation analyses. These data indicate that the frontal cortical contribution to SDAmC performance is anatomically selective: PL, but not IL or lOFC, is necessary to modulate behavior under motivational conflict.

**Figure 3.**
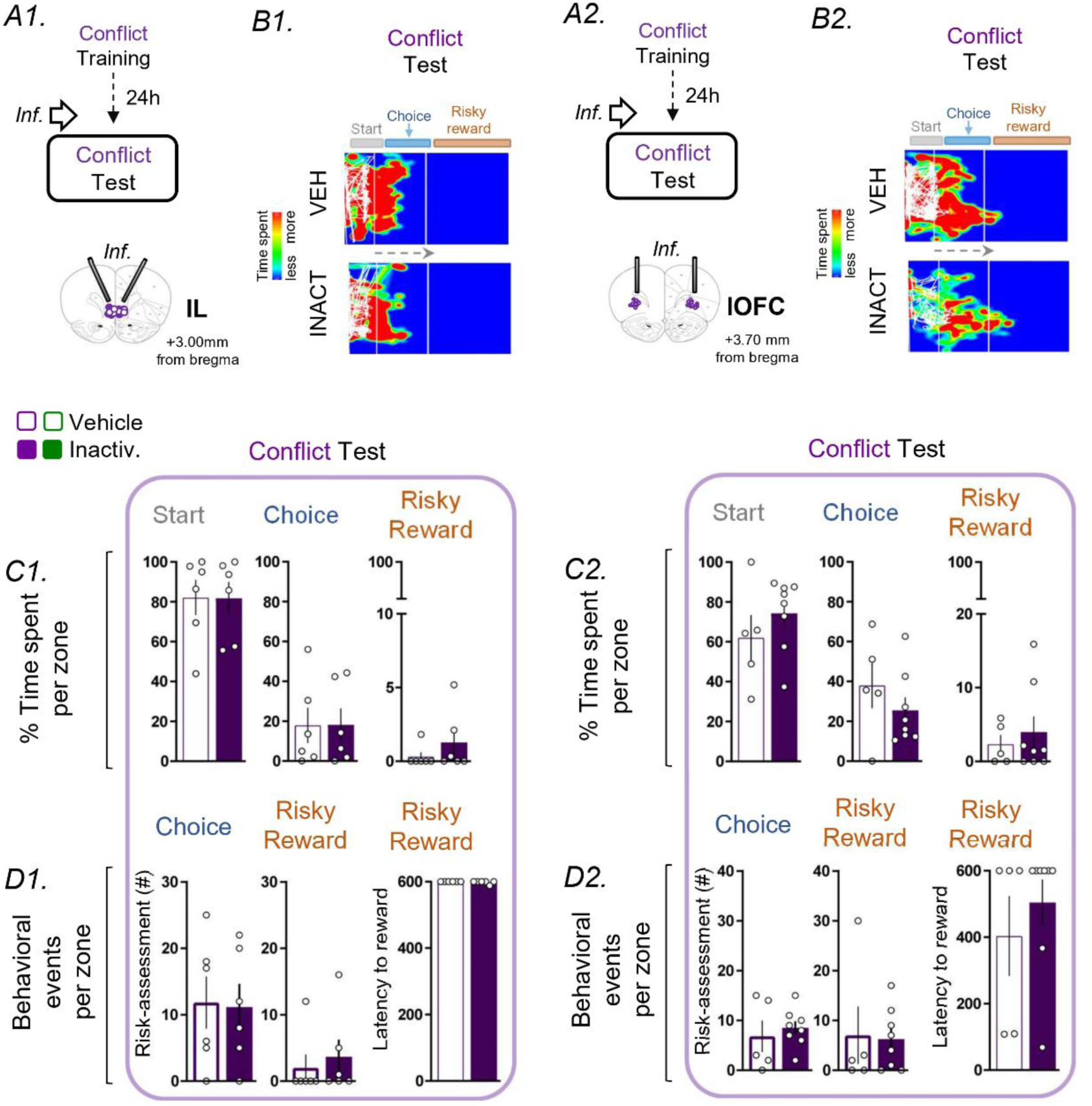
Inactivation of infralimbic cortex (IL) or lateral orbitofrontal cortex (lOFC) does not alter conflict behavior. (A1–A2) Cannula placements and timelines for IL (A1) and lOFC (A2) inactivation prior to Conflict testing. (B1–B2) Representative heatmaps showing comparable spatial occupancy patterns under vehicle (VEH) and inactivation (INACT) conditions. (C1–C2) Time spent in the start, choice, and reward zones during Conflict for IL (C1) and lOFC (C2), showing no significant differences. (D1–D2) Risk assessment measures and step-down latency remained unchanged following IL or lOFC inactivation. IL: (vehicle, n = 6; inactivation, n = 6); lOFC: (vehicle, n = 5; inactivation, n = 8). Data are shown as mean ± SEM. Individual data points are represented by open circles.

### Posterior insular cortex inactivation facilitates conflict resolution

The posterior insular cortex (pIC; cannula placements shown in **Fig. 4A**) was targeted based on its role in processing interoceptive state and aversive consequences (Gehrlach et al., 2019; Livneh et al., 2017). Representative heatmaps illustrate increased occupancy of grid zones following pIC inactivation during conflict, with no obvious change under non-conflict conditions (**Fig. 4B**). During conflict, vehicle-treated rats showed the expected avoidance-biased pattern (**Fig. 4B, top**). pIC inactivation facilitated entry onto the grid while diminishing active risk sampling at the decision point: step-down latency was markedly reduced (Mann–Whitney, U = 4.5, p < 0.001; **Fig. 4D**), and risk assessment events in the choice zone decreased (Mann–Whitney, U = 16.5, p = 0.010; **Fig. 4D**), whereas time in start zone did not differ significantly (p > 0.05; **Fig. 4C**). These effects were selective to motivational conflict: in a separate cohort tested under non-conflict conditions, pIC inactivation did not alter behavior (all p > 0.05; **Fig. 4E–F**). Correlation analyses indicated reorganization of behavioral structure. Under vehicle, time in start was negatively associated with choice risk assessment (rs = −0.70, p = 0.035), and reward time was strongly coupled to risk assessment (rs = 0.96, p < 0.001). After pIC inactivation, the start–risk assessment association was lost (rs = 0.26, p = 0.445), reward time–risk assessment coupling weakened (rs = 0.45, p = 0.168), and a strong positive association emerged between time in start and step-down latency (rs = 0.84, p = 0.001), absent under vehicle conditions (**Fig. 4G**). Saccharin intake was unchanged (p > 0.05; **Fig. 4H**). Together, these results identify pIC as a critical node that constrains approach during conflict, with effects dissociable from PL by virtue of reduced risk assessment.

**Figure 4.**
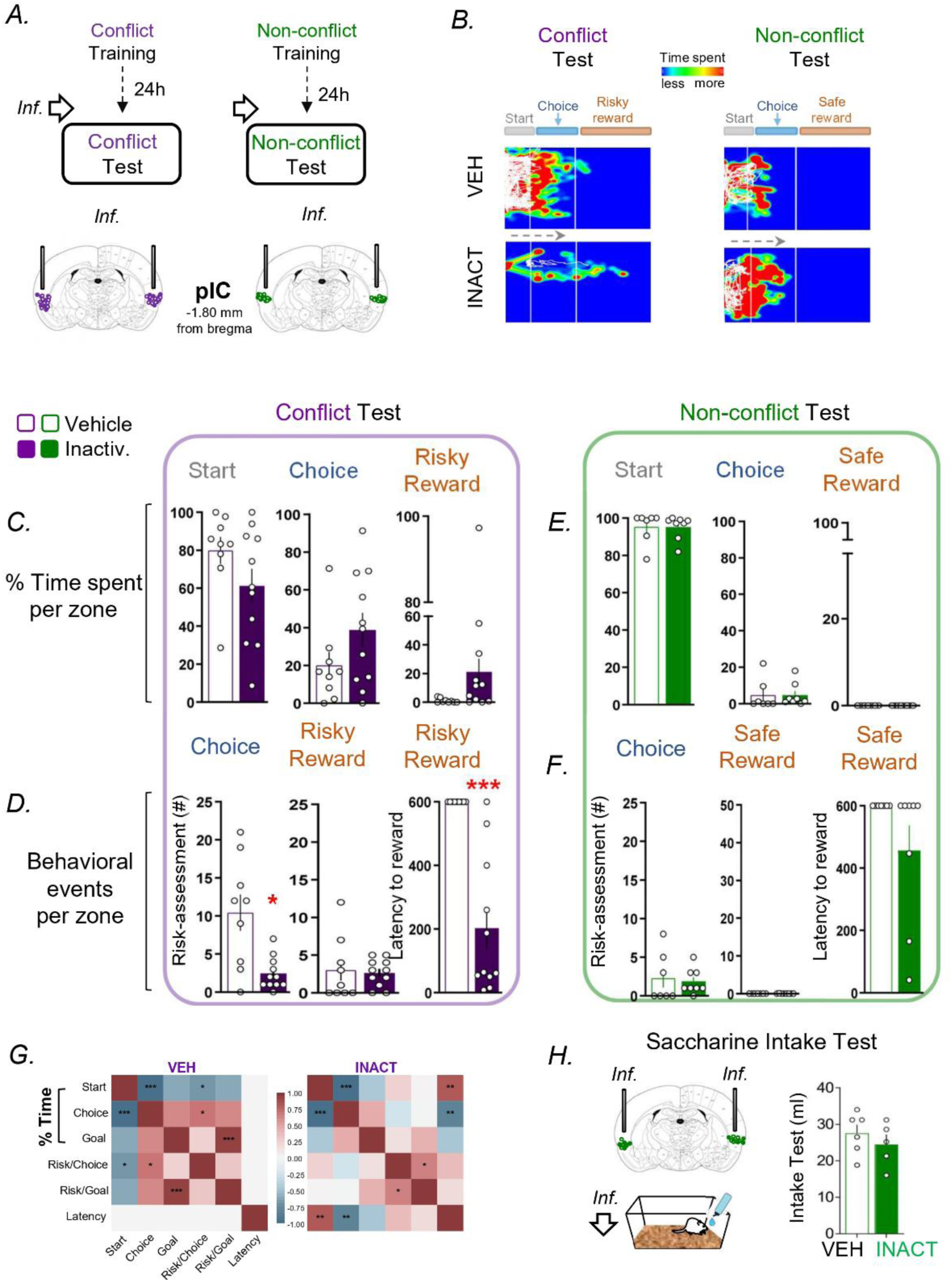
Posterior insular cortex (pIC) inactivation facilitates conflict resolution and reduces risk assessment. (A) Cannula placements for pIC and timeline indicating microinfusions prior to Conflict or Non-conflict testing. (B) Representative heatmaps illustrating behavior during Conflict and Non-conflict sessions under vehicle (VEH) or inactivation (INACT). (C–D) Conflict test quantification showing that pIC inactivation reduced step-down latency and decreased risk assessment in the choice zone, with no significant change in time in the start zone. (E and F) Non-conflict test measures showing no detectable effects of pIC inactivation under satiety. (G) Correlation matrices illustrating behavioral organization during Conflict under VEH versus INACT conditions. (H) Home-cage saccharin intake showing no effect of pIC inactivation on consummatory behavior (vehicle, n = 6; inactivation, n = 5). Conflict group (vehicle, n = 9; inactivation, n = 11); non-conflict group (vehicle, n = 7; inactivation, n = 8). *p < 0.05; **p < 0.01; ***p < 0.001. Data are expressed as mean ± SEM. Individual data points are represented by open circles.

### Inactivation of aIC and LHb does not alter conflict behavior

To further define anatomical specificity, we tested the anterior insular cortex (aIC; **Fig. 5A1**), implicated in salience-related control of approach (Rogers-Carter, Djerdjaj, Gribbons, Varela, & Christianson, 2019), and the lateral habenula (LHb; **Fig. 5A2**), implicated in aversive signaling and avoidance bias (Shabel et al., 2014; Stamatakis & Stuber, 2012; Velazquez-Hernandez & Sotres-Bayon, 2021). Representative heatmaps show no apparent shift in spatial distribution following aIC or LHb inactivation during conflict (**Fig. 5B1–B2**). Consistent with this, inactivation of either region did not alter conflict behavior: spatial allocation across the start, choice, and reward zones, risk assessment frequency, and step-down latency were comparable between vehicle and inactivation conditions (all p > 0.05; **Fig. 5C1–C2, D1–D2**). Given the absence of primary effects during conflict, these regions were not evaluated further. These null results constrain the network identified here, highlighting an anterior-posterior dissociation within insula cortex (pIC, but not aIC) and indicating that LHb is not required for approach-avoidance competition in this paradigm.

**Figure 5.**
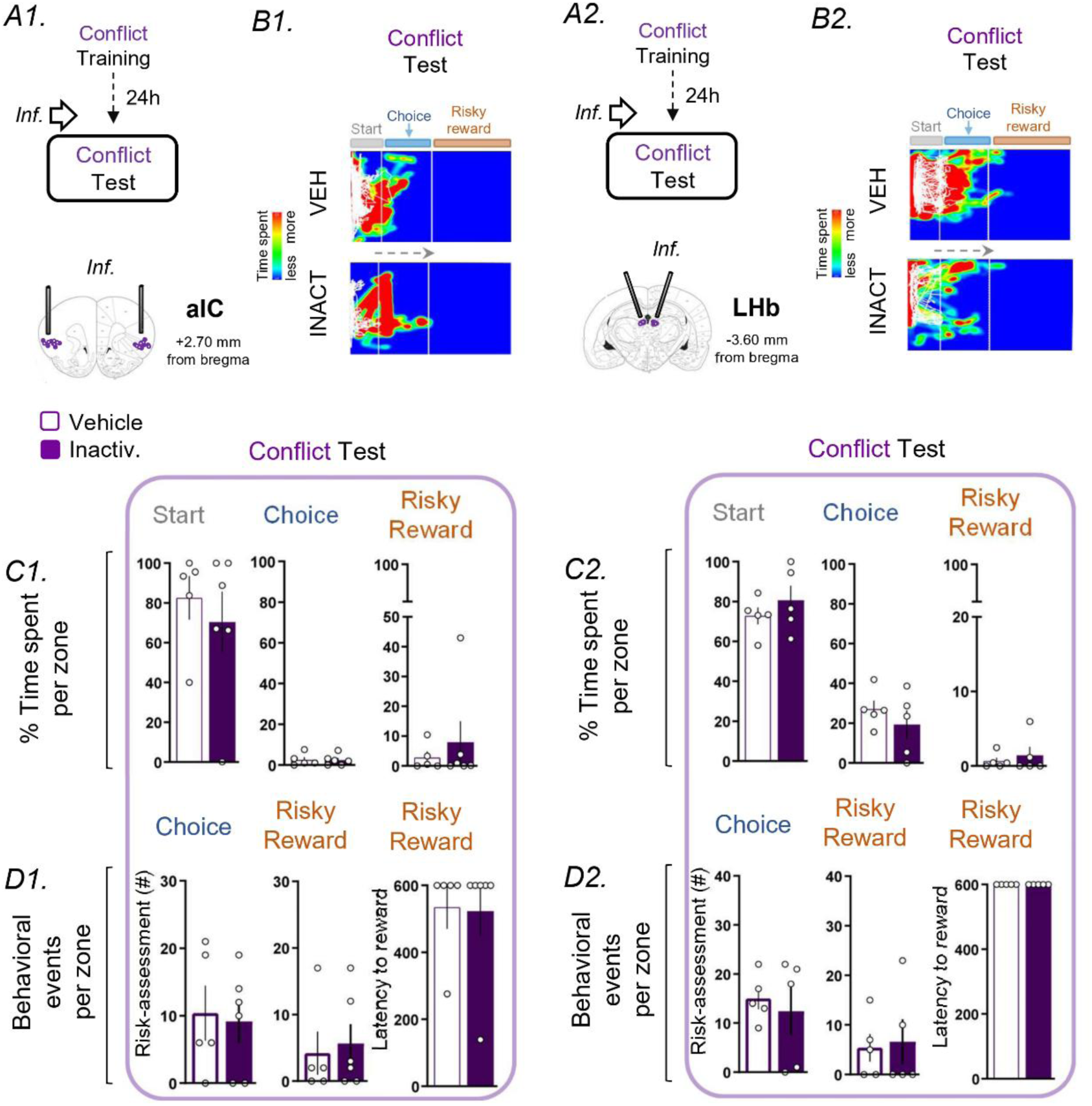
Inactivation of anterior insular cortex (aIC) or lateral habenula (LHb) does not alter conflict behavior. (A1–A2) Cannula placements and timelines for aIC (A1) and LHb (A2) inactivation prior to Conflict testing. (B1–B2) Representative heatmaps showing similar spatial occupancy under vehicle (VEH) and inactivation (INACT) conditions. (C1–C2) Time spent in the start, choice, and reward zones during Conflict for aIC (C1) and LHb (C2), showing no significant differences. (D1–D2) Risk assessment measures and step-down latency remained unchanged following aIC or LHb inactivation. aIC: (vehicle, n = 5; inactivation, n = 5); LHb: (vehicle, n = 5; inactivation, n = 5). Data are shown as mean ± SEM. Individual data points are represented by open circles.

### Basolateral amygdala inactivation induces generalized loss of avoidance

The basolateral amygdala (BLA; cannula placements shown in **Fig. 6A**) was targeted given its central role in valence encoding and threat memory expression (Beyeler et al., 2016; Blair, Sotres-Bayon, Moita, & Ledoux, 2005; Janak & Tye, 2015; Namburi et al., 2015). Representative heatmaps illustrate increased occupancy of grid and reward-proximal zones following BLA inactivation during both conflict and non-conflict testing (**Fig. 6B**). During conflict, BLA inactivation reduced time in the start zone (Mann–Whitney, U = 13.00, p = 0.043; Fig. 6C), decreased risk assessment in choice (Mann–Whitney, U = 14.50, p = 0.047; **Fig. 6D**), and reduced step-down latency (Mann–Whitney, U = 9.00, p = 0.008; **Fig. 6D**), consistent with facilitated approach. Unlike PL and pIC, BLA inactivation also altered behavior in the non-conflict test: in satiated rats, time in the reward zone increased (Mann–Whitney, U = 3.00, p = 0.030; **Fig. 6E**) and step-down latency decreased (Mann–Whitney, U = 2.00, p = 0.015; **Fig. 6F**), suggesting generalized loss of avoidance independent of motivational state. Correlation analyses supported a reorganization of behavioral structure. Under vehicle, time in the choice zone was perfectly correlated with risk assessment in reward (rs = 1.00, p < 0.001), whereas after BLA inactivation, this relationship reversed (rs = −0.86, p = 0.029). Risk assessment events in choice and reward became perfectly coupled (rs = 1.00, p < 0.001) and were strongly associated with step-down latency (rs = 0.85, p = 0.034), indicating altered coordination of risk sampling dynamics (**Fig. 6G**). Overall, BLA integrity is necessary for avoidance expression in this task, and its inactivation produces generalized facilitation of approach across motivational states.

**Figure 6.**
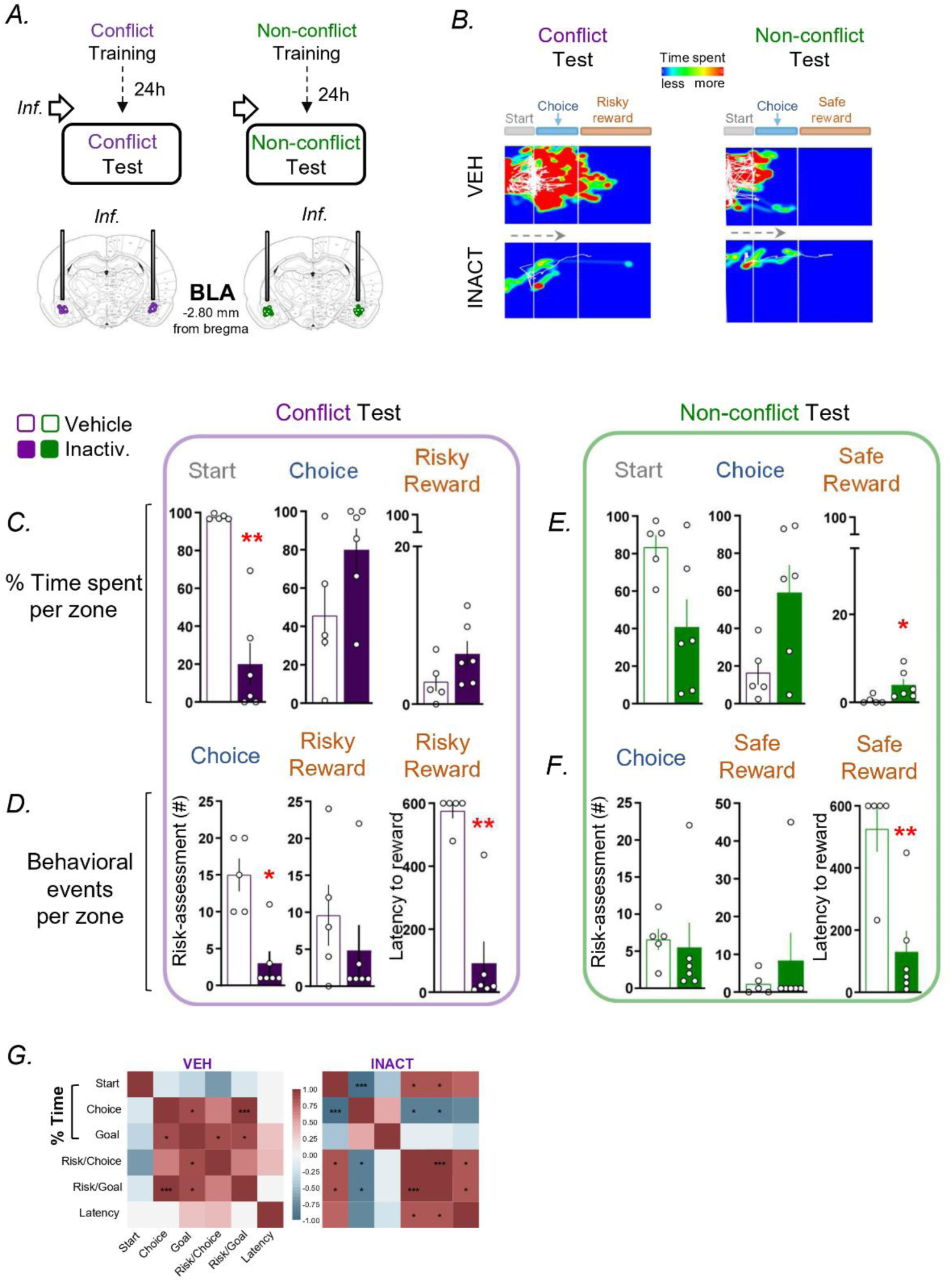
Basolateral amygdala (BLA) inactivation induces generalized loss of avoidance across motivational states. (A) Cannula placements for BLA and timeline indicating microinfusions prior to Conflict or Non-conflict testing. (B) Representative heatmaps showing increased progression onto the grid following inactivation (INACT) during both conflict and non-conflict sessions. (C–D) Conflict test measures showing reduced time in start, reduced risk assessment in choice, and reduced step-down latency after BLA inactivation. (E–F) Non-conflict test measures showing that BLA inactivation facilitated approach even under satiety, including increased time in reward and reduced step-down latency. (G) Correlation matrices illustrating behavioral organization during Conflict under vehicle (VEH) versus inactivation (INACT) conditions. Conflict group (vehicle, n = 5; inactivation, n = 6); non-conflict group (vehicle, n = 5; inactivation, n = 6). *p < 0.05; **p < 0.01; ***p < 0.001. Data are shown as mean ± SEM. Individual data points are represented by open circles.

### Nucleus accumbens inactivation induces a disorganized approach response

The nucleus accumbens (NAc; cannula placements shown in **Fig. 7A**) was targeted given its role in translating motivational state into goal-directed action (Ambroggi, Ishikawa, Fields, & Nicola, 2008; Floresco, 2015; Salamone & Correa, 2012; Salamone et al., 2016). Representative heatmaps show increased progression toward the grid and reward zones after NAc inactivation during conflict, with minimal change in non-conflict conditions (**Fig. 7B**). During conflict, NAc inactivation reduced time in the start zone (Mann–Whitney, U = 7, p = 0.042; **Fig. 7C**) and increased time in the choice and reward zones (Mann–Whitney, U = 7, p = 0.042; and U = 4.5, p = 0.012; **Fig. 7C**), while reducing step-down latency (Mann–Whitney, U = 4, p = 0.011; **Fig. 7D**). However, this progression toward the goal was accompanied by a disruption in risk assessment organization: risk assessment decreased in choice (Mann–Whitney, U = 6, p = 0.026; **Fig. 7D**) but increased in reward (Mann–Whitney, U = 7.5, p = 0.043; **Fig. 7D**), consistent with impaired coordination of risk sampling across spatial zones. In the non-conflict test, NAc inactivation did not alter spatial distribution or step-down latency (all p > 0.05; Fig. **7E–F**), and saccharin intake was unaffected (**Fig. 7H**), arguing against nonspecific motor impairment or reduced motivation. Correlation analyses showed reorganization: under vehicle, reward time was perfectly correlated with reward risk assessment (rs = 1.00, p < 0.001), whereas after NAc inactivation this association weakened (rs = 0.60, p = 0.088). Time in the start zone became strongly negatively correlated with time in the reward zone (rs = −0.83, p = 0.005) and positively correlated with step-down latency (rs = 0.85, p = 0.004); time in the choice and reward zones became positively correlated (rs = 0.83, p = 0.005), and time in the reward zone became negatively associated with step-down latency (rs = −0.83, p = 0.005) (**Fig. 7G**). Together, these results indicate that NAc integrity is required to maintain the normal coupling between spatial progression, risk sampling, and action timing during motivational conflict.

**Figure 7.**
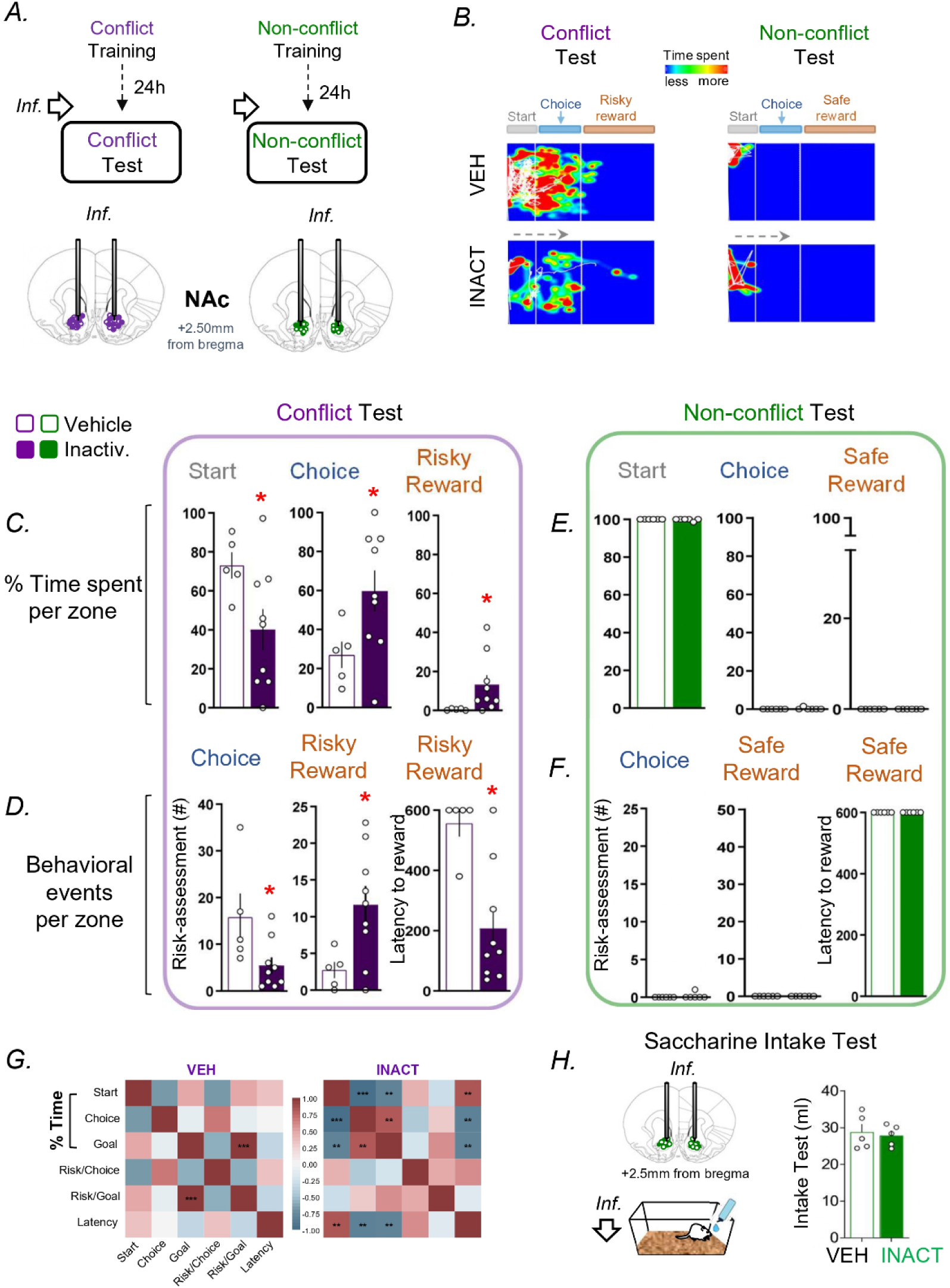
Nucleus accumbens (NAc) inactivation increases progression toward the goal while disrupting risk assessment organization during conflict. (A) Cannula placements for NAc and timeline indicating microinfusions prior to Conflict or Non-conflict testing. (B) Representative heatmaps showing increased occupancy on the grid following inactivation (INACT) during Conflict, with minimal change under non-conflict conditions. (C–D) Conflict test measures showing reduced time in the start zone, increased time in the choice and reward zones, reduced step-down latency, and a mixed redistribution of risk assessment (decreased in the choice zone, increased in reward). (E–F) Non-conflict test measures showing no detectable effects of NAc inactivation under satiety. (G) Correlation matrices illustrating behavioral organization during Conflict under vehicle (VEH) versus inactivation (INACT) conditions. (H) Home-cage saccharin intake showing no effect of NAc inactivation on consummatory behavior (vehicle, n = 5; inactivation, n = 5). Conflict group (vehicle, n = 5; inactivation, n = 9); non-conflict group (vehicle, n = 6; inactivation, n = 6). *p < 0.05; **p < 0.01; ***p < 0.001. Data are shown as mean ± SEM. Individual data points are represented by open circles.

Across structures, these results reveal a constrained and anatomically selective network supporting conflict behavior. PL and pIC inactivation biased performance toward approach specifically under motivational conflict, but with dissociable impacts on risk assessment, whereas BLA inactivation produced a generalized loss of avoidance evident even in non-conflict conditions. In contrast, NAc inactivation promoted spatial progression toward the goal while disrupting the coordination of risk assessment across zones, consistent with impaired organization of goal-directed action under threat. Finally, several candidate regions commonly implicated in valuation or aversive signalling (IL, lOFC, aIC, and LHb) were dispensable in this paradigm. We next integrate these positive and negative findings to identify the minimal set of hubs required to organize avoidance, risk assessment, and approach during reward seeking under threat.

## Discussion

In this study, we used a validated approach-avoidance paradigm (Illescas-Huerta et al., 2021) and systematic pharmacological inactivation to identify which brain regions are necessary to organize behavior during reward-seeking under threat. Our main finding is that conflict performance depends on a constrained cortico-limbic network, rather than a broad engagement of regions commonly implicated in threat or valuation. Among the eight candidate structures tested, only PL, pIC, BLA, and NAc showed detectable effects on conflict behavior, and each region contributed in a dissociable manner. PL and pIC inactivation facilitated approach selectively under motivational conflict, BLA inactivation produced a generalized loss of avoidance that extended to non-conflict conditions, and NAc inactivation disrupted the coordination of risk assessment and approach despite increasing progression toward the reward. In contrast, IL, lOFC, aIC, and LHb did not show detectable effects under the present experimental conditions, providing an informative set of negative results that constrains mechanistic models of approach-avoidance conflict. Together, these findings support a selective architecture in which a limited set of hubs is necessary to organize avoidance, risk assessment, and approach when reward pursuit competes with threat. These coupling patterns reflect functional covariation rather than anatomical connectivity. A consolidated view of the positive and null effects is provided in Table 1, while a network-level interpretation of the critical hubs is provided in Fig. 8A-B.

**Figure 8.**
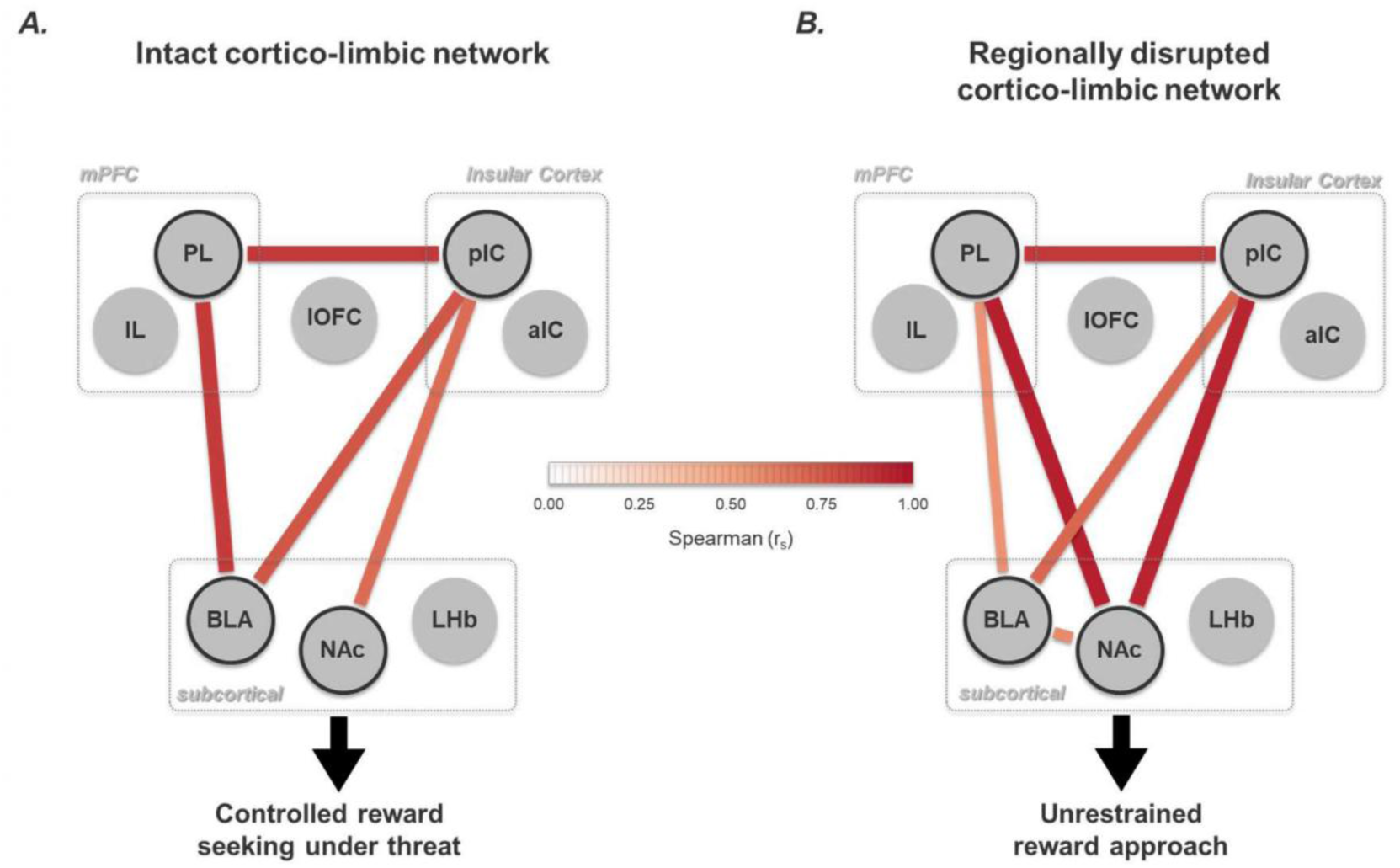
Functional network schematic comparing the vehicle and regional inactivation conditions. Schematic representation of the cross-regional functional coupling among the main nodes implicated by the SDAmC, shown separately for the vehicle condition (A) and the regional inactivation condition (B). Circles denote regional nodes (PL, IL, lOFC, pIC, aIC, BLA, NAc, LHb), dashed boxes group nodes into cortical networks. Black-outlined nodes highlight regions identified as critical hubs in the behavioral analyses (i.e., regions whose inactivation produced robust changes in the conflict phenotype), whereas non-outlined/grey nodes indicate regions without a detectable behavioral impact under the present paradigm. Edges depict Spearman rank correlations (rs) between nodes (color coded-bar) and represent statistical associations rather than anatomical connectivity. This schematic summarizes the task-specific network configuration supporting conflict-related avoidance, risk assessment, and the organization of goal-directed action, and illustrates how regional inactivation reshapes coupling patterns among the key hubs.

**Table 1.** Summary of behavioral effects of regional inactivation in the SDAmC. Table summarizing the direction and specificity of behavioral changes produced by regional inactivation relative to vehicle across the SDAmC. For each region, effects are indicated separately for the conflict and non-conflict conditions and across the three task zones (start, choice, reward), as well as the main derived measures (risk assessment at choice and reward, and latency). Upward (green) and downward (red) arrows denote significant increases or decreases relative to the vehicle, respectively. “–” indicates no detectable effect under the present paradigm, whereas Fig.” indicates that the behavioral variable was not measurable because there was no initial conflict-related effect during the evaluation.

| Summary of inactivation results |  |  |  |  |  |  |  |  |  |  |  |  |
| --- | --- | --- | --- | --- | --- | --- | --- | --- | --- | --- | --- | --- |
| Brain region | Conflict |  |  |  |  |  | Non-conflict |  |  |  |  |  |
|  | Time in Start | Time in Choice | Time in Goal | Risk Assess. in Choice | Risk Assess. in Goal | Latency | Time in Start | Time in Choice | Time in Goal | Risk Assess. in Choice | Risk Assess. in Goal | Latency |
| PL | ↓ | ↑ | — | — | — | ↓ | — | — | — | — | — | — |
| IL | — | — | — | — | — | — | n.a. | n.a. | n.a. | n.a. | n.a. | n.a. |
| IOFC | — | — | — | — | — | — | n.a. | n.a. | n.a. | n.a. | n.a. | n.a. |
| pIC | — | — | — | ↓ | — | ↓ | — | — | — | — | — | — |
| aIC | — | — | — | — | — | — | n.a. | n.a. | n.a. | n.a. | n.a. | n.a. |
| LHb | — | — | — | — | — | — | n.a. | n.a. | n.a. | n.a. | n.a. | n.a. |
| BLA | ↓ | — | — | ↓ | — | ↓ | — | — | ↑ | — | — | ↓ |
| NAc | ↓ | ↑ | ↑ | ↓ | ↑ | ↓ | — | — | — | — | — | — |

### PL gates conflict-specific approach without increasing overt risk sampling

PL inactivation biased conflict resolution toward approach, reflected by reduced time in the start zone, increased time in the choice zone, and a strong reduction in step-down latency. These effects were selective to conflict conditions, suggesting that PL is preferentially recruited when threat and reward compete. This is consistent with convergent evidence implicating PL in defensive control, persistent monitoring of threat, and the expression of avoidance strategies (Bravo-Rivera et al., 2014; Sierra-Mercado et al., 2011; Sotres-Bayon & Quirk, 2010; Sotres-Bayon et al., 2012). Importantly, PL inactivation did not increase the frequency of risk assessment, indicating that the approach bias was not accompanied by heightened overt sampling of the grid. Instead, correlation analyses showed that PL inactivation reorganized the internal structure of behavior, particularly by decoupling risk assessment across zones and altering relationships between start occupancy, risk assessment, and latency. This pattern suggests PL primarily maintains a coherent defensive strategy linking evaluation to action timing under motivational competition, rather than generating risk assessment events per se. Within the SDAmC framework, PL may therefore act as a cortical hub that stabilizes a conservative strategy, preserving hesitation and threat sensitivity when appetitive drive is high.

### pIC constrains approach by supporting decision-point risk evaluation

pIC inactivation also facilitated approach under conflict, but with a distinct signature: a large reduction in step-down latency accompanied by decreased risk assessment in the choice zone. These effects were absent in non-conflict conditions and occurred without changes in home-cage saccharin intake, arguing against nonspecific changes in motivation. The pIC has been implicated in processing aversive states and interoceptive signals that shape avoidance and threat-related valuation (Gehrlach et al., 2019; Livneh et al., 2017). In this task, pIC appears particularly important at the decision point, where animals typically probe the grid and sample threat proximity before committing to descent. The loss of risk assessment after pIC inactivation suggests that this region supports a component of conflict behavior that is not reducible to avoidance output alone but rather reflects active evaluation under uncertainty. Notably, the dissociation between aIC and pIC is consistent with the idea that insula is functionally heterogeneous (Gehrlach et al., 2019; Livneh et al., 2017; T. Shi, Feng, Wei, & Zhou, 2020; W. Shi, Fu, Shi, & Zhou, 2022), and that posterior subdivisions may be more directly coupled to aversive consequences and bodily state computations relevant for approach-avoidance arbitration.

Importantly, the effects of PL and pIC inactivation are unlikely to reflect nonspecific locomotor activation or generalized increases in reward motivation. These inactivations did not alter behavior under non-conflict conditions and did not affect saccharin consumption, indicating that these manipulations selectively altered conflict-dependent action selection.

### BLA supports avoidance across motivational states

BLA inactivation produced a qualitatively different phenotype from PL or pIC: facilitated approach during conflict was accompanied by similar facilitation in non-conflict conditions, including increased time near the reward and reduced step-down latency under satiety. This generalized effect is consistent with BLA’s central role in threat memory expression and associative control of defensive behavior (Bravo-Rivera et al., 2014; Burgos-Robles et al., 2017; Hernandez-Jaramillo, Illescas-Huerta, & Sotres-Bayon, 2024; Namburi et al., 2015; Pickens et al., 2003; Tye et al., 2011). In SDAmC, the grid carries an instrumental threat contingency acquired during threat conditioning; weakening that association would be expected to reduce avoidance regardless of motivational state. Thus, while PL and pIC modulated how behavior is organized under competition, BLA integrity appeared necessary for maintaining the aversive constraint that defines the task. This interpretation also aligns with extensive work positioning the amygdala as a key hub for linking learned threat to behavioral output, which can then be shaped by prefrontal regulation depending on context and competing motivations (Dallerac et al., 2017; Moscarello & LeDoux, 2013; Pape & Pare, 2010; Sierra-Mercado et al., 2011; Sotres-Bayon & Quirk, 2010).

### NAc coordinates coherent goal-directed action under threat

NAc inactivation increased progression toward the reward under conflict, reducing time in the start zone and decreasing step-down latency, but produced a distinctive disorganization of risk assessment: risk assessment decreased at the choice zone yet increased at the reward zone. This mixed pattern suggests a disruption in the temporal organization of exploration and evaluation, rather than simple facilitation of approach. NAc is classically involved in translating motivational state into action selection and vigor (Ambroggi et al., 2008; Stuber et al., 2011). Within SDAmC, conflict requires not only initiating movement toward the reward but coordinating when and where risk sampling occurs as the animal advances across threat-associated space. The reorganization of behavioral correlations after NAc inactivation supports the view that NAc contributes to coupling spatial progression with structured evaluation, enabling goal-directed action to unfold coherently under threat. That NAc inactivation did not alter non-conflict behavior or saccharin intake further argues against a primary deficit in locomotion or appetite, pointing instead to a role in organizing the structure of conflict-specific action.

### Informative null effects refine the candidate network for motivational conflict

A central contribution of this study is the systematic inclusion of regions that were plausible candidates but produced no measurable behavioral changes in this task. IL, often associated with suppression of defensive responding and extinction-related processes (Sierra-Mercado et al., 2011; Vidal-Gonzalez et al., 2006), did not alter conflict performance, suggesting that IL is not required for organizing the avoidance-risk assessment-approach sequence measured here. Likewise, lOFC did not affect conflict behavior despite its established involvement in updating action strategies and encoding outcome information (Gremel & Costa, 2013; Orsini et al., 2015). One possibility is that lOFC becomes more relevant under conditions requiring flexible updating, contingency shifts, or probabilistic outcomes, whereas SDAmC conflict performance depends more strongly on stable arbitration between an appetitive goal and a learned threat contingency. Similarly, aIC inactivation produced no detectable effects, supporting regional specificity within the insular cortex, and suggesting that anterior insular cortex contributions may be more prominent in tasks emphasizing salience, social valuation, or complex affective appraisal (Rogers-Carter et al., 2019). Finally, LHb inactivation did not alter conflict behavior despite its prominent role in aversive signaling and avoidance bias (Shabel et al., 2014; Stamatakis & Stuber, 2012). LHb may be more critical under conditions involving punishment prediction errors, uncontrollable stressors, or broader state-dependent negative bias, whereas the instrumental avoidance demands of SDAmC may rely more directly on amygdala-dependent aversive memory and prefrontal/insular regulation of action selection.

Importantly, these null effects should not be interpreted as evidence that these regions are irrelevant for motivated decision-making in general, but rather that they are not necessary for the specific components of conflict behavior quantified in SDAmC under the present conditions. By incorporating negative results alongside positive ones, the current dataset provides a sharper constraint on circuit models of motivated conflict and helps distinguish conflict-specific hubs from regions whose contributions may emerge under different task demands.

### Limitations and future directions

Several factors frame the interpretation of these findings. First, experiments were conducted in adult male rats, and sex-dependent mechanisms may modulate conflict processing; this should be addressed given increasing evidence for sex differences in defensive regulation and motivated behavior (Day, Suwansawang, Halliday, & Stevenson, 2020; Gruene, Roberts, Thomas, Ronzio, & Shansky, 2015; Orsini & Setlow, 2017; Orsini, Willis, Gilbert, Bizon, & Setlow, 2016). Second, muscimol/baclofen inactivation provides strong causal inference at the regional level but does not resolve projection-specific mechanisms; future work using pathway-selective manipulations will be necessary to determine whether the PL–NAc, pIC–striatal, or amygdala–striatal axes contribute differentially to the behavioral signatures observed here. Third, the SDAmC task captures a specific conflict configuration in which appetitive drive is manipulated by thirst and the threat contingency is learned instrumentally. Future studies varying motivational state, threat controllability, or reinforcement uncertainty may reveal additional region-specific contributions, including potential roles for structures such as the LHb or OFC under more dynamic decision contexts.

## Conclusions

Together, these results identify a minimal cortico-limbic architecture to organize reward seeking under threat. PL and pIC constrain conflict-specific approach through dissociable effects on the coordination of risk assessment and action timing, BLA maintains the aversive constraint necessary for avoidance across motivational states, and NAc supports coherent organization of goal-directed progression under conflict. By systematically testing both candidate and non-candidate regions within the same behavioral framework, this work provides causal evidence that motivated conflict behavior is supported by a selective network of hubs rather than a diffuse “emotional” system, offering a clearer foundation for circuit-level models of approach-avoidance arbitration.

## Author Contributions

ROS, LR-L, EI-H and FS-B designed research; ROS, LR-L, EI-H and AR-B performed experiments; ROS, LR-L, EI-H, AR-B and FS-B analyzed data; and ROS, EI-H and FS-B wrote the paper.

## Acknowledgements

We thank Jorge A. Quillfeldt for providing the original step-down avoidance chamber, Rafael Villafuerte Peralta for assistance during early stages of behavioral data collection, and members of the Sotres Lab for helpful discussions and comments on the manuscript.

